# Lizards in energy-saving mode: Urban individuals are smaller and have lower metabolic rates

**DOI:** 10.1101/2025.02.02.635261

**Authors:** Rebecca Rimbach, Öncü Maraci, Avery L. Maune, Isabel Damas-Moreira

**Affiliations:** Department of Behavioural Biology, University of Münster; Joint Institute for Individualisation in a Changing Environment (JICE), University of Münster and Bielefeld University, Münster, Germany; Department of Behavioural Ecology, Bielefeld University

**Keywords:** energy expenditure, HIREC, Lacertidae, metabolism, physiology, *Podarcis muralis*, scaled mass index, urbanization

## Abstract

Human-induced rapid environmental change is intensifying globally. Urbanization, a major form of human-induced rapid environmental change, alters ecosystems and exposes wildlife to urban stressors, including altered ambient temperatures and resource shifts. These alterations can impose selective pressures on wildlife, influencing behavior, physiology, and morphology. Alterations in resource availability and quality, and ambient temperature may be especially important for ectotherms, which depend on environmental conditions for regulating body temperature and activity. To date, little research has examined the physiological changes of ectotherms that are associated with urban life. In this study, we measured standard metabolic rate (SMR; N = 38) and recorded body mass and size in common wall lizards (*Podarcis muralis*) from an urban (N = 123) and a semi-natural habitat (N = 125) in Croatia. Our results show that urban lizards exhibit a lower SMR, reduced body mass, and shorter snout-vent length than their semi-natural counterparts. Despite these differences, body condition, assessed via the scaled mass index, did not vary between the two habitat types, potentially reflecting adaptive responses to urbanization-related environmental pressures. These findings suggest adaptive strategies of reptiles in urban environments and highlight the importance of studying both physiological and morphological traits to better understand the effects of urbanization on wildlife.

## Introduction

Urbanization represents one of the most prominent human-induced rapid environmental changes (HIREC) to the natural environment (McKinney, 2002; Sih et al., 2011). It simultaneously alters several ecological conditions, exposing wildlife to different types of stressors, such as pollution (e.g., light, noise and air), altered resource availability and quality, novel and altered biotic interactions with pathogens, competitors and predators (Alberti et al., 2020; Isaksson and Bonier, 2020; Sanders et al., 2021). Furthermore, urban structures and human activities lead to the ‘urban heat island effect’, in which urban areas experience higher temperatures than surrounding rural areas, changing the microclimatic conditions in urban habitats (Imhoff et al., 2010). As urban environments keep expanding worldwide, it is imperative to study the species that can live in urban areas and the extent to which they are affected by altered environmental conditions in urban habitats. The first adjustment to cope with altered environments is often behavioral (Tuomainen and Candolin, 2011). Many urban populations behave differently from their non-urban counterparts (Gaynor et al., 2018; Lowry et al., 2013; Ritzel and Gallo, 2020; Sih et al., 2011; Sol et al., 2013; Thompson et al., 2018). The additive effects of intrinsic (e.g. behavioral) and extrinsic (e.g. environmental) changes can have profound impacts on physiology and morphology, and ultimately fitness consequences. Yet, how urban environments drive physiological and morphological changes has received little attention (Alberti, 2015; Bonier, 2012; French et al., 2018; Putman and Tippie, 2020).

Maintenance metabolism represents the minimal cost of living and is a major functional trait because it restricts the allocation of energy to survival, growth, and reproduction (Glazier, 2015). Maintenance metabolism is defined as the minimal energy expenditure of a post-absorptive resting endotherm or ectotherm, and is measured as resting (RMR) or standard metabolic rate (SMR), respectively. Across different taxa, environmental changes associated with urbanization and global warming have been found to alter metabolic rates. For instance, urban populations of Eurasian red squirrels (*Sciurus vulgaris*) display weaker metabolic responses to changes in ambient temperatures compared to their rural counterparts (Wist et al., 2023). Urban greater white-toothed shrews (*Crocidura russula*) and great tits (*Parus major*) exhibit lower RMRs than their rural counterparts (Andersson et al., 2018; Oliveira et al., 2020). While traffic noise decreases the RMR of house sparrow (*Passer domesticus*) nestlings (Brischoux et al., 2017), human presence at night does not alter RMR of house finches (*Haemorhous mexicanus*) from urban and rural habitats (Hutton et al., 2018). These examples demonstrate that metabolic responses to urbanization-related environmental changes are complex and can be species-specific or dependent on the type of environmental stressor, highlighting the need for additional studies on a variety of taxa before more general conclusions can be drawn. Changes in metabolic rates in response to urban conditions may have implications for organisms by impacting survival and reproduction, directly influencing their fitness (Norin and Metcalfe, 2019; Pettersen et al., 2018).

Ectotherms are likely to be affected by urbanization differently than endotherms because they are highly dependent on environmental temperatures and habitat characteristics to regulate their body temperature, with profound consequences for behavior and physiology (Angilletta, 2009; Stevenson, 1985). Thus, studying the metabolic adjustments of ectotherms in response to environmental change, such as urbanization and global warming, is crucial for predicting population persistence and species resilience (Nowakowski et al., 2018). However, research on the effects of urbanization on metabolic changes in ectotherms has been limited to date, with mixed results. Urban acorn-dwelling ants (*Temnothorax curvispinosus*) exhibit higher metabolic rates than rural ants when tested at 25°C (Chick et al., 2021), as do common toads (*Bufo bufo*) exposed to artificial light at night (Touzot et al., 2019). In contrast, rock agamas (*Laudakia vulgaris*) and Mediterranean house geckos (*Hemidactylus turcicus*) from urban and natural habitats do not differ in SMR (Vardi et al., 2023). Despite these insights, several questions remain to better understand or predict how ectotherms that often inhabit cities, such as several lizard species, respond physiologically to anthropogenic habitat alterations (Doherty et al., 2020; French et al., 2018; Putman and Tippie, 2020). Several biogeographic comparative studies report that ectotherms from warmer climates have lower metabolic rates compared to those from cooler climates (Addo-Bediako et al., 2002; DeLong et al., 2018; Gaston et al., 2009). This countergradient variation has been interpreted as an adaptation to cold climates characterized by short growing seasons. Recently, this view has also been applied to discussions of climate change and warming temperatures, where reduced metabolic rates may help mitigate the negative effects of heat stress (Pilakouta et al., 2020). Similar processes and adjustments may also underlie the reduction of metabolic rates observed in some urban populations, potentially due to the ‘urban heat island effect’ (Imhoff et al., 2010).

Similarly, body size — a trait closely linked to metabolism — also shows diverse patterns in response to urbanization. A global meta-analysis found that lizards tend to have larger body sizes, and a similar body condition, in urban habitats compared to more natural ones (Putman and Tippie, 2020). However, these findings are based on geographically and taxonomically biased data (i.e., studies from the United States and Australia contributed more than 66% of effect sizes and *Anolis* lizards provided 75% of effect sizes), limiting its generalizability across clades. For example, rock agamas (*L. vulgaris*), house geckos (*H. turcicus*) and side-blotched lizards (*Uta stansburiana*) exhibit similar body size and mass across urban and natural habitats (Putman et al., 2024; Vardi et al., 2023). In contrast, urban Western fence lizards (*Sceloporus occidentalis*) and Australian water dragons (*Intellagama lesueurii*) exhibit a smaller body size compared to individuals from more natural habitats (Putman et al., 2024). These mixed responses show, similarly to studies on metabolic rates within an urban ecology framework, that more research is needed to better understand the observed patterns across lizard species.

The Lacertidae lizards in Europe are globally among the most affected reptile families due to anthropogenic habitat modification (Doherty et al., 2020). Yet, this family also includes a few species with a remarkable ability to thrive in anthropogenic habitats, such as the common wall lizard (*Podarcis muralis*). This species is very successful in urbanized habitats, both along its native and invasive range (Ferner, 2004; Williams, 2019), and is often found in city centers, exposed to high levels of habitat modification, artificial structures, and anthropogenic disturbance. However, little is known about its physiological and morphological adjustments to urban life. Urban common wall lizards exhibit higher developmental instability compared to rural populations, measured by fluctuating asymmetry in morphological traits (Lazić et al., 2013), indicating that urban populations may experience greater environmental stress. To better understand the physiological and morphological adjustments of ectotherms to urbanization, the goals of this study were to assess differences in i) metabolic rates (measured as SMR), ii) body size and mass, and iii) body condition between common wall lizards from urban and semi-natural habitats. Given that urban habitats expose animals to altered environmental conditions, such as elevated temperatures, reduced resource availability, and increased pollution and anthropogenic disturbance, we hypothesized that urban lizards would exhibit lower metabolic rates compared to those from semi-natural habitats. This could reflect an energy-saving strategy to cope with the challenges posed by urban environments. Given the available, albeit biased, data on urbanization-related changes in reptile body size (Putman and Tippie, 2020), we predicted that urban individuals would be larger compared to lizards from semi-natural areas. If urban lizards must conserve energy to cope with the challenges of the urban environment, we expect them to have a lower body condition compared to individuals living in semi-natural habitats.

## Methods

### Study sites and animals

We studied 38 common wall lizards (*Podarcis muralis*), living in two different sites of Rovinj, Croatia in June 2024. We captured 19 individuals (9 females, 9 males) in the city center (45.082639, 13.636167), where the level of habitat modification, artificial structures, and anthropogenic disturbance is high. At the second site, a semi-natural area (45.065344, 13.652087), we captured 19 individuals (7 females, 12 males). This area was located 3 km from the city center, and consists of a conifer forest, predominantly composed of natural substrate, with low amounts of artificial structures, and with some degree of human disturbance during the day.

We captured animals with a noose and transported lizards in an individually numbered breathable linen bag. To ensure a post-absorptive state before SMR measurements (Mell et al., 2016; Žagar et al., 2015), we kept lizards individually for 2 days in small terraria (18 cm x 33 cm, with a height of 13 cm), at a room temperature of 25 °C and without providing any food. Each terrarium was equipped with a shelter and a small ceramic bowl with *ad libitum* water. Daily, we placed a hand warmer (Firebag™; warms up to 50 °C) next to the shelter area to allow lizards to thermoregulate (Damas-Moreira et al., 2024).

### Measurements of standard metabolic rate

We measured SMR using an open-circuit system with a Field Metabolic System (FMS, Sable Systems, Las Vegas NV, USA) between 6:50 am and 2:15 pm. We weighed individuals using a digital scale (± 0.1g) and then placed one individual in an air-tight metabolic chamber (55 mm diameter, 680 ml nominal volume). Subsequently, we initiated measurements and video recording of the behavior of individuals in the metabolic chambers using an action camera (CU-SPC06, Cooau, Shenzhen, China). We measured SMR at 25°C (± 0.61 °C). Readings of O_2_ and CO_2_ were taken every second for 4 x 10 min during a total of one hour of measurement. Before each 10-min window, the baseline chamber was referenced for 5 min. Air was pulled through the system at a rate of 100 ml/min. Six to seven animals were measured per day, on average 3.8 urban and 3.2 non-urban individuals per day; the order of individuals on any given day was chosen randomly. After the measurements, we weighed individuals again and used average body mass for analysis. We processed metabolic records using a macro program recorded in ExpeData software (Sable Systems). We calculated SMR from the lowest level of O_2_ consumption recorded for 90 consecutive readings (i.e. 90 sec). We confirmed the resting state of individuals by visual inspection of oxygen consumption rates and video recordings of lizards inside the chamber. All data were corrected for drift in O_2_, CO_2_, and H_2_O baselines using the Drift Correction function in ExpeData.

### Morphometric measurements

After metabolic rate measurements, we measured each lizard’s snout-to-vent length (SVL) to the nearest 0.1 mm using a digital caliper. All measurements of SVL were conducted by a single researcher (A.L.M.). Subsequently, we released individuals at the site of capture. In addition to these 38 individuals, we also measured body size and mass for an additional 210 common wall lizards (80 females, 130 males) captured in the city Rovinj and the same semi-natural habitat in the years 2021, 2022 and 2023 during previous studies (data collected in 2021 has been published in Damas-Moreira et al., 2024). Body mass and SVL were determined by two researchers (I.D.M & A.L.M.), using the same methods as described above. This resulted in a total sample size of 248 lizards (semi-natural: N = 125; urban: N = 123) for which we measured body mass and SVL. For all individuals, we calculated the scaled mass index (SMI) from the regression of log-transformed SVL and log-transformed body mass. SMI accounts for the allometric relationship between body mass and body length and is a standardized measure of body condition (Peig and Green, 2009).

### Statistical analysis

We conducted all analyses using R version 4.4.1 (R Core Team, 2024). To assess habitat-specific differences in SMR (N = 38), we fitted a linear mixed model (LMM) using the lme4 package (Bates et al., 2015). We included SMR as the response variable and habitat type (urban vs. semi-natural), sex, average body mass, temperature during the SMR measurement and time of day (in minutes) as fixed effects, and cycle number as a random factor.

For the large dataset (N = 248), we used a linear model (LM) to assess habitat differences in body mass because a LMM including year as a random factor resulted in a singular fit. We used LMMs to assess habitat differences in snout-vent length (SVL) and body condition (SMI), and included habitat type (urban vs. semi-natural) and sex as fixed effects and year as a random factor. We tested for one-way interactions between the fixed effects habitat type and sex.

For all models, we determined the contribution of interaction terms using likelihood ratio tests (LRT) by comparing the model with the interaction term to the model without. We excluded the interaction term because the explanatory power of the model was not improved by the inclusion of the interaction based on the LRT (Pinheiro and Bates, 2000). For all models, we checked that model assumptions were met using the functions ‘check_model’, ‘check_singularity’ and ‘check_normality’ in the performance package (Lüdecke et al., 2021). We plotted predictor effects plots using the effects package (Fox and Weisberg, 2018; Fox and Weisberg, 2019) and the ggplot2 package (Wickham, 2016).

## Results

### Standard metabolic rate

Urban lizards tended to have a lower SMR (1.26 ± 0.54 kJ/day) than lizards from the semi-natural habitat (1.64 ± 0.70 kJ/day; Fig. 1A; Table 1). Males had a higher SMR (1.70 ± 0.64 kJ/day) compared to females (1.06 ± 0.43 kJ/day; Fig. 1B; Table 1).

**Figure 1.**
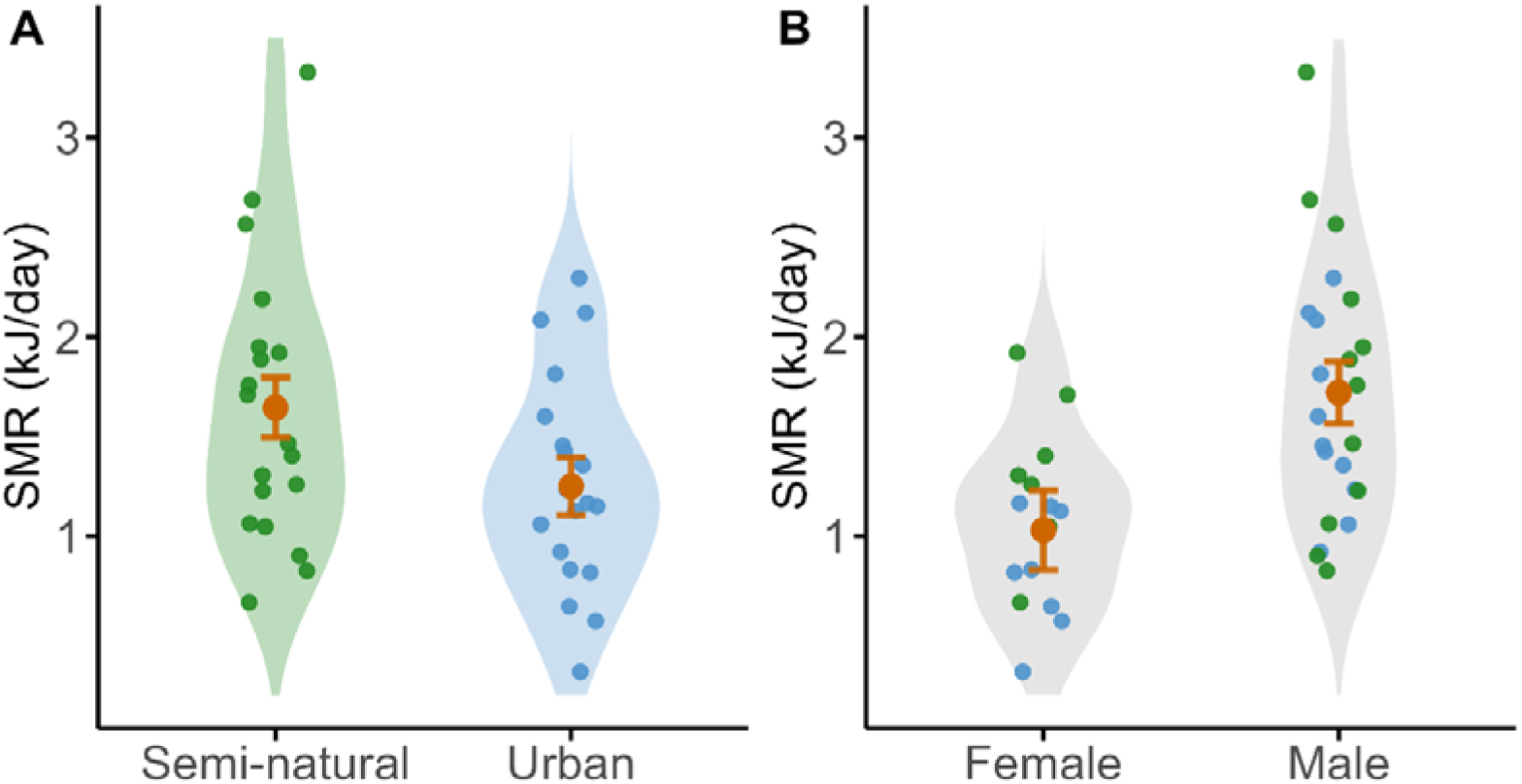
Habitat and sex differences in standard metabolic rate (SMR) of common wall lizards (*Podarcis muralis*). (A) Urban individuals tend to have a lower SMR than individuals from a semi-natural habitat (LMM: p = 0.057). (B) Female lizards have lower SMR than males (LMM: p = 0.019). Green points indicate raw data from lizards of the semi-natural habitat (N = 19), blue points data from urban lizards (N = 19), and orange points and error bars (SE) predictions from the LMM. The shaded areas represent kernel density plots.

**Table 1.** Results of a linear mixed model to test for habitat differences in standard metabolic rate (SMR) of adult common wall lizards (*Podarcis muralis*; N = 38). The models included habitat type (urban vs. semi-natural), sex, average body mass, temperature during the SMR measurement, and time of day as explanatory variables, and cycle number as a random factor (reference categories are indicated in []).

| *Predictors* | **Effects on SMR (kJ/day)** |  |  |
| --- | --- | --- | --- |
|  | *Estimates* | *CI* | *p* |
| (Intercept) | -1.70 | -10.88 – 7.48 | 0.708 |
| Habitat [Urban] | -0.39 | -0.80 – 0.01 | 0.057 |
| Sex [Male] | 0.69 | 0.12 – 1.26 | 0.019 |
| Average body mass | -0.02 | -0.23 – 0.19 | 0.824 |
| Temperature | 0.14 | -0.24 – 0.53 | 0.447 |
| Time of day | -0.06 | -0.15 – 0.04 | 0.236 |
| **Random Effects** |  |  |  |
| $\sigma^2$ | 0.306 | | |
| $\tau_{00 \text{ cycle}}$ | 0.010 | | |
| ICC | 0.032 |  |  |
| N<sub>cycle</sub> | 5 |  |  |
| Observations | 38 |  |  |
| Marginal R<sup>2</sup> / Conditional R<sup>2</sup> | 0.306 / 0.328 |  |  |

Note: The marginal R<sup>2</sup> considers only the variance of the fixed effects, while the conditional R<sup>2</sup> takes both the fixed and random effects into account. Abbreviations: $\sigma^2$ , variance of the random effect “year”; $\tau_{00}$ , variance of the random intercept at a specific grouping level; ICC, Intraclass coefficient of variation.

### Morphometrics and body condition

The larger dataset also showed that urban lizards (4.55 ± 1.21 g, N = 123) had a smaller body mass than lizards from the semi-natural habitat (4.88 ± 1.25 g, N = 125; LM: Estimate = -0.28, CI = -0.55 – -0.02, p = 0.036; Fig. 2A), and males were heavier (5.21 ± 1.18 g, N = 153) than females (3.93 ± 0.86, N = 95; LM: Estimate = 1.27, CI = 1.00 – 1.54, p = <0.001; adjusted R^2^ = 0.263; Fig. 2B). Urban lizards also had a smaller SVL (60.4 ± 5.16 mm; Fig. 2C) than lizards from the semi-natural habitat (62.0 ± 4.76 mm), and males had a larger SVL (62.0 ± 5.13 mm) than females (59.9 ± 4.56 mm; Fig. 2D; Table 2). SMI did not differ between habitat types (urban: 4.68 ± 0.62 g, N = 123; semi-natural: 4.61 ± 0.68 g, N = 125; Fig. 2E), but males had a higher SMI (4.93 ± 0.55 g, N = 153) compared to females (4.19 ± 0.52 g, N = 95; Fig. 2F; Table 2).

**Figure 2.**
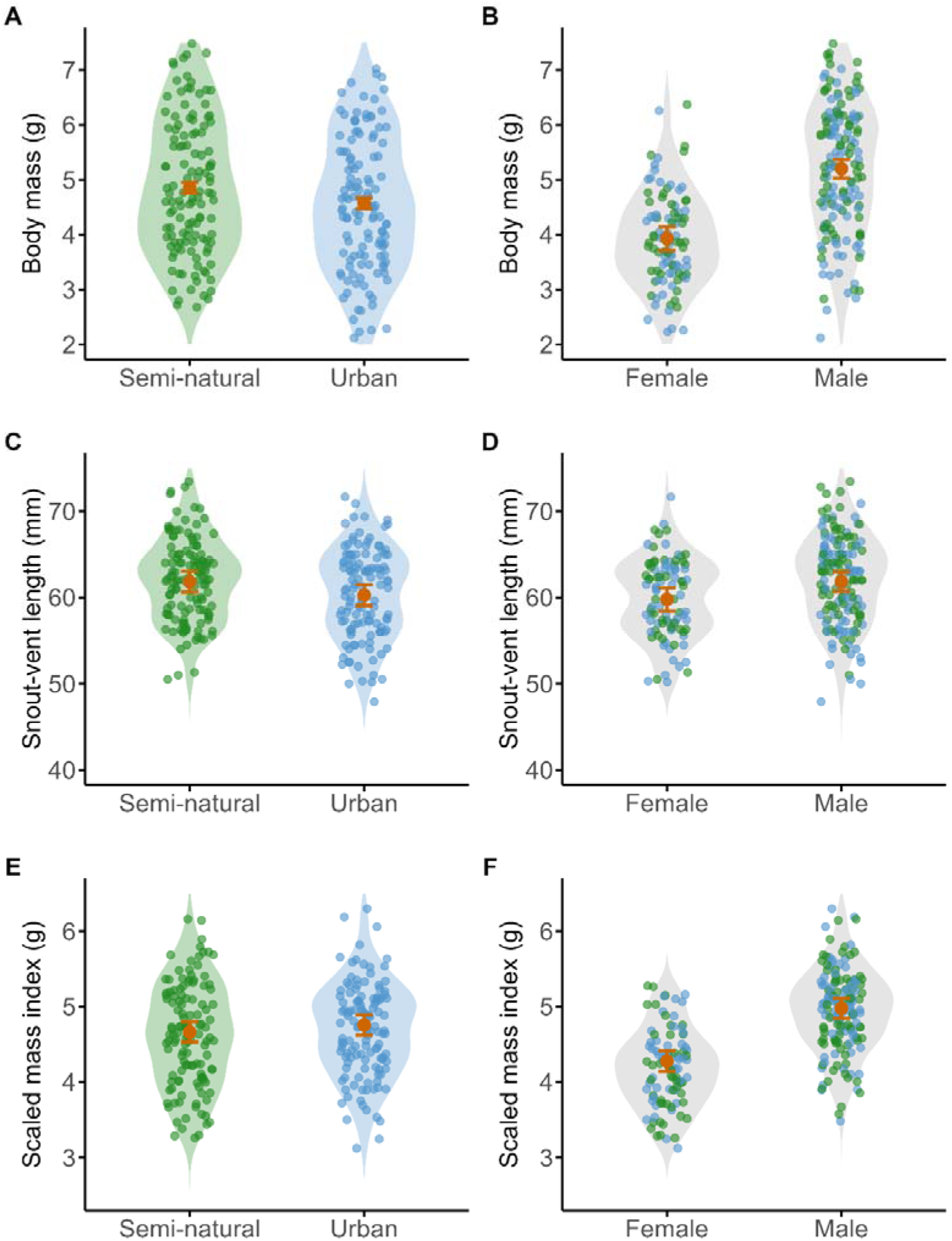
Habitat and sex differences in body mass, snout-vent length and scaled mass index in common wall lizards (*Podarcis muralis*). (A) Urban common wall lizards (*Podarcis muralis*) have a lower body mass (LM: p = 0.036) and (C) a smaller snout-vent length (LMM: p = 0.012), but (E) do not differ in scaled mass index from lizards from a semi-natural habitat (LMM: p = 0.148).

**Table 2.** Results of linear mixed models to test for habitat differences in snout-vent length (SVL) and scaled mass index (SMI) of adult common wall lizards (*Podarcis muralis*; N = 248). The models included habitat type (urban vs. semi-natural) and sex as explanatory variables, and year as a random factor (reference categories are indicated in []).

| *Predictors* | **Effects on SVL** |  |  | **Effects on SMI** |  |  |
| --- | --- | --- | --- | --- | --- | --- |
|  | *Estimates* | *CI* | *p* | *Estimates* | *CI* | *p* |
| (Intercept) | 60.54 | 59.07 – 62.01 | <0.001 | 4.23 | 3.95 – 4.51 | <0.001 |
| Habitat [Urban] | -1.56 | -2.76 – -0.35 | 0.012 | 0.09 | -0.03 – 0.22 | 0.148 |
| Sex [Male] | 2.09 | 0.79 – 3.39 | 0.002 | 0.70 | 0.56 – 0.84 | <0.001 |
| **Random Effects** |  |  |  |  |  |  |
| $\sigma^2$ | 23.218 | | | 0.249 | | |
| $\tau_{00 \text{ year}}$ | 0.708 | | | 0.064 | | |
| ICC | 0.030 |  |  | 0.204 |  |  |
| $N_{\text{year}}$ | 4 | | | 4 | | |
| Observations | 248 |  |  | 248 |  |  |
| Marginal $R^2$ | 0.066 | | | 0.273 | | |
| Conditional $R^2$ | 0.094 | | | 0.421 | | |

Note: The marginal $R^2$ considers only the variance of the fixed effects, while the conditional $R^2$ takes both the fixed and random effects into account. Abbreviations: $\sigma^2$ , variance of the random effect “year”; $\tau_{00}$ , variance of the random intercept at a specific grouping level; ICC, Intraclass coefficient of variation.

Males are heavier (LM: p = <0.001), larger (LMM: p = 0.002) and have a higher scaled mass index than females (LM: p < 0.001; B, D, F). Green points indicate raw data from lizards of the semi-natural habitat (N = 125), blue points data from urban lizards (N = 123), and orange points and error bars (SE) predictions from the linear (mixed) models. Shaded area represents kernel density plots.

## Discussion

Our study investigated the physiological and morphological differences between common wall lizards inhabiting an urban and a semi-natural habitat. We found that urban lizards exhibited a lower SMR compared to their semi-natural conspecifics, along with smaller body size (SVL) and mass. We did not detect differences in body condition (SMI) between habitat types. These findings highlight physiological and morphological adjustments in common wall lizards to urban environments and contribute to our understanding of how ectotherms might adjust to anthropogenic habitat modification.

Additionally, we found that males had a higher SMR than females, consistent with findings in other ectotherms (Angilletta and Sears, 2000; Niewiarowski and Waldschmidt, 1992; but see, for example, Orrell et al., 2004). This difference in metabolic rate may reflect differences in behavior, physiological demands, or reproductive strategies between sexes. Future research could investigate sex-specific energetic mitigation strategies for urban life and examine how sex-specific behaviors and environmental pressures drive metabolic differences.

Our results show a trend for urban lizards to have a lower SMR than lizards from the semi-natural habitat, supporting our initial hypothesis and indicating that urban individuals may downregulate their metabolism in response to urban stressors. Lower metabolic rates have also been observed in endotherms coping with urban-related environments (Andersson et al., 2018; Oliveira et al., 2020). This reduction in metabolic rate could represent an energy-saving strategy to cope with altered environmental conditions in urban areas, such as elevated temperatures, reduced resource availability, and the need to mitigate additional urban stressors, such as pollution and anthropogenic disturbance. Reduced SMR could minimize energy expenditure when resources are limited or of lower quality, helping individuals maintain energy balance in urban habitats. Exposure to environmental stressors or elevated glucocorticoid levels can alter metabolic rates and mitochondrial function (Norin and Metcalfe, 2019; Voituron et al., 2022), and result in a downregulation of SMR (in toad-headed agamas, *Phrynocephalus przewalskii*; Han et al., 2023). It would be important to further understand whether a metabolic downregulation could be an adaptive strategy for wall lizards to cope with the unique challenges of urban environments.

Contrary to our predictions, urban lizards had both smaller body size and mass compared to lizards from semi-natural habitats. This contrasts with previous studies that found larger body sizes in urban reptiles (Putman and Tippie, 2020). However, smaller body sizes in urban lizards have also been reported in some species (Baxter-Gilbert et al., 2021; Putman et al., 2024). This may result from several factors, including altered resource availability or quality, increased exposure to pollutants or differences in thermal resources, such as access to radiant heat (Mugabo et al., 2010; Sinervo and Adolph, 1989). Individuals that experience stressors during development, i.e. developmental stress, can show altered growth patterns and reduced body size and condition, as demonstrated in zebra finches (Kraft et al., 2019). Moreover, some studies report high lizard population densities in urban areas, potentially affecting individual access to resources (Mugabo et al., 2010). Smaller body sizes could also reflect evolutionary pressures, as urban environments may favor individuals with faster life histories or lower energy demands (Brans and De Meester, 2018). Additionally, if urban habitats consist of smaller anthropogenic crevices, smaller body sizes could be favored in cities, allowing lizards to shelter and fit into tighter spaces. The smaller body sizes we observed in urban wall lizards might also have implications for other fitness-related traits, such as locomotor performance, predator escape, or reproductive success (Tatu et al., 2024). Future research should investigate the ecological and evolutionary implications of these morphological changes.

Despite differences in body size and mass, we found no difference in body condition between urban and semi-natural lizards, suggesting that body condition is maintained across habitats. This may indicate that urban lizards are capable of adjusting their behavior and/or physiology and morphology to optimize resource use despite the challenges posed by urban environments. For instance, urban lizards could adjust their daily activity patterns to minimize energy expenditure or exploit food sources available around anthropogenic structures, such as insects attracted by artificial night at light. Interestingly, Indian rock agamas (*Psammophilus dorsalis*) also exhibit no difference in body condition between urban and rural habitats, but urbanization triggered physiological coping responses, including lower heterophil-to-lymphocyte ratios and lower testosterone levels (Amdekar et al., 2018). Our findings add to the growing evidence on the impact of urbanization on urban populations and that ectotherms exhibit diverse and context-dependent responses to urbanization. While lower metabolic rates and smaller body sizes may represent adaptive responses to the challenges of urban environments, these adjustments may also have long-term consequences for fitness and population dynamics. For instance, reduced body size may limit reproductive output, while a lower SMR could constrain the ability to respond to additional environmental stressors. Future studies, including replicates of urban and non-urban habitats, temporal replication, and larger sample sizes, will provide further insight into the metabolic responses of lizards and other ectotherms to urbanization-related environmental change.

The variation in responses to urbanization observed among reptile species highlights the need for more taxonomically and geographically diverse studies. The differences in SMR and body size between urban and semi-natural common wall lizards underscore the importance of studying physiological and morphological traits together to understand how organisms adjust to urbanization. Further research could investigate a potential genetic basis of these changes, the role of developmental plasticity, and the interaction between urban stressors such as pollution, habitat structure, and temperature.

## Conclusions

Overall, our results demonstrate that common wall lizards exhibit habitat-specific physiological and morphological adjustments to urbanization. Urban lizards have lower SMR and smaller body sizes than lizards from a semi-natural habitat, suggesting that these traits may reflect compensatory responses to the altered environmental conditions in urban areas. These findings suggest adaptive strategies of reptiles in urban environments and highlight the importance of studying both physiological and morphological traits to better understand the effects of urbanization on wildlife. As urbanization continues to expand globally, further research on the mechanisms underlying these responses will be crucial for predicting species resilience and informing conservation strategies.

## Declarations

### Authors’ contributions

R.R., Ö.M., and I.D.M. designed the study; R.R., A.L.M. and I.D.M. collected all data; R.R. conducted the statistical analyses. R.R. conceptualized the manuscript and wrote the initial draft. All authors (R.R., Ö.M., A.L.M., and I.D.M.) assisted with drafting and editing the manuscript.

## Acknowledgements

We thank Duje Lisičić and Tobias Wittenbreder for all their help and support with field work. We also thank Melanie Dammhahn for commenting on a previous version of this manuscript.

## Ethics approval

Lizard captures and experimental procedures were approved by the Department for Environmental and Nature Protection of the Croatian Ministry of Economy and Sustainable Development, under the permit number 517-10-1-2-24-2 of the class UP/l-352-04/24-08/80.

## References

Addo-Bediako, A., Chown, S. L. and Gaston, K. J. (2002). Metabolic cold adaptation in insects: a large-scale perspective. Funct. Ecol. 16, 332–338.

Alberti, M. (2015). Eco-evolutionary dynamics in an urbanizing planet. Trends Ecol. Evol. 30, 114–126.

Alberti, M., Palkovacs, E. P., Des Roches, S., De Meester, L., Brans, K. I., Govaert, L., Grimm, N. B., Harris, N. C., Hendry, A. P., Schell, C. J., et al. (2020). The complexity of urban eco-evolutionary dynamics. Bioscience 70, 772–793.

Amdekar, M. S., Kakkar, A. and Thaker, M. (2018). Measures of health provide insights into the coping strategies of urban lizards. Front. Ecol. Evol. 6, 128.

Andersson, M. N., Nilsson, J., Nilsson, J. Å. and Isaksson, C. (2018). Diet and ambient temperature interact to shape plasma fatty acid composition, basal metabolic rate and oxidative stress in great tits. J. Exp. Biol. 221, jeb.186759.

Angilletta, M. J. (2009). Thermal adaptation: a theoretical and empirical synthesis. London: Oxford University Press.

Angilletta, M. J. and Sears, M. W. (2000). The metabolic cost of reproduction in an oviparous lizard. Funct. Ecol. 14, 39–45.

Bates, D., Mächler, M., Bolker, B. and S, W. (2015). Fitting linear mixed-effects models using lme4. J. Stat. Softw. 67, 1–48.

Baxter-Gilbert, J., Riley, J. L., Frère, C. H. and Whiting, M. J. (2021). Shrinking into the big city: influence of genetic and environmental factors on urban dragon lizard morphology and performance capacity. Urban Ecosyst. 24, 661–674.

Bonier, F. (2012). Hormones in the city: endocrine ecology of urban birds. Horm. Behav. 61, 763–772.

Brans, K. I. and De Meester, L. (2018). City life on fast lanes: Urbanization induces an evolutionary shift towards a faster lifestyle in the water flea *Daphnia*. Funct. Ecol. 32, 2225–2240.

Brischoux, F., Meillère, A., Dupoué, A., Lourdais, O. and Angelier, F. (2017). Traffic noise decreases nestlings’ metabolic rates in an urban exploiter. J. Avian Biol. 48, 905–909.

Chick, L. D., Waters, J. S. and Diamond, S. E. (2021). Pedal to the metal: cities power evolutionary divergence by accelerating metabolic rate and locomotor performance. Evol. Appl. 14, 36–52.

Damas-Moreira, I., Szabo, B., Drosopoulos, G., Stober, C., Lisičić, D. and Caspers, B. A. (2024). Smarter in the city? Lizards from urban and semi-natural habitats do not differ in a cognitive task in two syntopic species. Curr. Zool. 70, 361–370.

DeLong, J. P., Bachman, G., Gibert, J. P., Luhring, T. M., Montooth, K. L., Neyer, A. and Reed, B. (2018). Habitat, latitude and body mass influence the temperature dependence of metabolic rate. Biol. Lett. 14, 20180442.

Doherty, T. S., Balouch, S., Bell, K., Burns, T. J., Feldman, A., Fist, C., Garvey, T. F., Jessop, T. S., Meiri, S. and Driscoll, D. A. (2020). Reptile responses to anthropogenic habitat modification: a global meta-analysis. Glob. Ecol. Biogeogr. 29, 1265–1279.

Ferner, J. (2004). The introduction of European and Italian wall lizards (*Podarcis muralis* and *P. sicula*; Reptilia, Lacertidae) into the United States. J. Ky. Acad. Sci. 65, 1–4.

Fox, J. and Weisberg, S. (2018). Visualizing fit and lack of fit in complex regression models with predictor effect plots and partial residuals. J. Stat. Softw. 87, 1–27.

Fox, J. and Weisberg, S. (2019). An R companion to applied regression. 3rd ed. Thousand Oaks CA: Sage Publications.

French, S. S., Webb, A. C., Hudson, S. B. and Virgin, E. E. (2018). Town and country reptiles: a review of reptilian responses to urbanization. Integr. Comp. Biol. 58, 948– 966.

Gaston, K. J., Chown, S. L., Calosi, P., Bernardo, J., Bilton, D. T., Clarke, A., Clusella-Trullas, S., Ghalambor, C. K., Konarzewski, M., Peck, L. S., et al. (2009). Macrophysiology: a conceptual reunification. Am. Nat. 174, 595–612.

Gaynor, K. M., Hojnowski, C. E., Carter, N. H. and Brashares, J. S. (2018). The influence of human disturbance on wildlife nocturnality. Science. 360, 1232–1235.

Glazier, D. S. (2015). Is metabolic rate a universal ‘pacemaker’ for biological processes? Biol. Rev. 90, 377–407.

Han, X., Sun, B., Zhang, Q., Teng, L., Zhang, F. and Liu, Z. (2023). Metabolic regulation reduces the oxidative damage of arid lizards in response to moderate heat events. Integr. Zool. 19, 1034–1046.

Hutton, P., Wright, C. D., DeNardo, D. F. and McGraw, K. J. (2018). No effect of human presence at night on disease, body mass, or metabolism in rural and urban house finches (*Haemorhous mexicanus*). Integr. Comp. Biol. 58, 977–985.

Imhoff, M. L., Zhang, P., Wolfe, R. E. and Bounoua, L. (2010). Remote sensing of the urban heat island effect across biomes in the continental USA. Remote Sens. Environ. 114, 504–513.

Isaksson, C. and Bonier, F. (2020). Urban evolutionary physiology. In Urban Evolutionary Biology (ed. Szulkin, M.), Munshi-South, J.), and Charmantier, A.), pp. 217–233. Oxford University Press.

Kraft, F. L. O. H., Driscoll, S. C., Buchanan, K. L. and Crino, O. L. (2019). Developmental stress reduces body condition across avian life-history stages: a comparison of quantitative magnetic resonance data and condition indices. Gen. Comp. Endocrinol. 272, 33–41.

Lazić, M. M., Kaliontzopoulou, A., Carretero, M. A. and Crnobrnja-Isailović, J. (2013). Lizards from urban areas are more asymmetric: using fluctuating asymmetry to evaluate environmental disturbance. PLoS One 8, e84190.

Lowry, H., Lill, A. and Wong, B. B. M. (2013). Behavioural responses of wildlife to urban environments. Biol. Rev. 88, 537–549.

Lüdecke, D., Ben-Shachar, M., Patil, I., Waggoner, P. and Makowski, D. (2021). performance: an R package for assessment, comparison and testing of statistical models. J. Open Source Softw. 60, 3139.

McKinney, M. L. (2002). Urbanization, biodiversity, and conservation. Bioscience 52, 883–890.

Mell, H., Josserand, R., Artacho, P. and Galliard, J. Le (2016). Do personalities co-vary with metabolic expenditure and glucocorticoid stress response in adult lizards? Behav. Ecol. Sociobiol. 70, 951–961.

Mugabo, M., Marquis, O., Perret, S. and Le Galliard, J. F. (2010). Immediate and delayed life history effects caused by food deprivation early in life in a short-lived lizard. J. Evol. Biol. 23, 1886–1898.

Niewiarowski, P. and Waldschmidt, S. R. (1992). Variation in metabolic rates of a lizard: use of SMAR in ecological contexts. Funct. Ecol. 6, 15–22.

Norin, T. and Metcalfe, N. B. (2019). Ecological and evolutionary consequences of metabolic rate plasticity in response to environmental change. Philos. Trans. R. Soc. B 374, 20180180.

Nowakowski, A. J., Watling, J. I., Thompson, M. E., Brusch, G. A., Catenazzi, A., Whitfield, S. M., Kurz, D. J., Suárez-Mayorga, Á., Aponte-Gutiérrez, A., Donnelly, M. A., et al. (2018). Thermal biology mediates responses of amphibians and reptiles to habitat modification. Ecol. Lett. 21, 345–355.

Oliveira, F. G., Mathias, M. da L., Rychlik, L., Tapisso, J. T. and von Merten, S. (2020). Metabolic and behavioral adaptations of greater white-toothed shrews to urban conditions. Behav. Ecol. 30, 1334–1343.

Orrell, K. S., Congdon, J. D., Jenssen, T. A., Michener, R. H. and Kunz, T. H. (2004). Intersexual differences in energy expenditure of *Anolis carolinensis* lizards during breeding and postbreeding seasons. Physiol. Biochem. Zool. 77, 50–64.

Peig, J. and Green, A. J. (2009). New perspectives for estimating body condition from mass/length data: the scaled mass index as an alternative method. Oikos 118, 1883– 1891.

Pettersen, A. K., Marshall, D. J. and White, C. R. (2018). Understanding variation in metabolic rate. J. Exp. Biol. 221, jeb166876.

Pilakouta, N., Killen, S. S., Kristjánsson, B. K., Skúlason, S., Lindström, J., Metcalfe, N. B. and Parsons, K. J. (2020). Multigenerational exposure to elevated temperatures leads to a reduction in standard metabolic rate in the wild. Funct. Ecol. 34, 1205–1214.

Pinheiro, J. C. and Bates, D. M. (2000). Mixed-effects models in S and S-Plus. New York, NY: Springer.

Putman, B. J. and Tippie, Z. A. (2020). Big city living: a global meta-analysis reveals positive impact of urbanization on body size in lizards. Front. Ecol. Evol. 8, 580745.

Putman, B. J., Rensel, M. A., Schlinger, B. A., French, S., Blumstein, D. T. and Pauly, G. B. (2024). Comparing fear responses of two lizard species across habitats varying in human impact. J. Urban Ecol. 10,.

R Core Team (2024). R: a language and environment for statistical computing. R Foundation for Statistical Computing, Vienna, Austria. www.r-project.org/.

Ritzel, K. and Gallo, T. (2020). Behavior change in urban mammals: a systematic review. Front. Ecol. Evol. 8, 576665.

Sanders, D., Frago, E., Kehoe, R., Patterson, C. and Gaston, K. J. (2021). A meta-analysis of biological impacts of artificial light at night. Nat. Ecol. Evol. 5, 74–81.

Sih, A., Ferrari, M. C. O. and Harris, D. J. (2011). Evolution and behavioural responses to human-induced rapid environmental change. Evol. Appl. 4, 367–387.

Sinervo, B. and Adolph, S. C. (1989). Thermal sensitivity of growth rate in hatchling *Sceloporus* lizards: environmental, behavioral and genetic aspects. Oecologia 78, 411–419.

Sol, D., Lapiedra, O. and González-Lagos, C. (2013). Behavioural adjustments for a life in the city. Anim. Behav. 85, 1101–1112.

Stevenson, R. D. (1985). The relative importance of behavioral and physiological adjustments controlling body temperature in terrestrial ectotherms. Am. Nat. 126, 362–386.

Tatu, A., Dutta, S. and Thaker, M. (2024). Hotter deserts and the impending challenges for the Spiny-tailed Lizard in India. Biol. Open 13, bio060150.

Thompson, M. J., Evans, J. C., Parsons, S. and Morand-Ferron, J. (2018). Urbanization and individual differences in exploration and plasticity. Behav. Ecol. 29, 1415–1425.

Touzot, M., Teulier, L., Lengagne, T., Secondi, J., Théry, M., Libourel, P. A., Guillard, L. and Mondy, N. (2019). Artificial light at night disturbs the activity and energy allocation of the common toad during the breeding period. Conserv. Physiol. 7, coz002.

Vardi, R., Dubiner, S., Bezalel, R. B., Meiri, S. and Levin, E. (2023). Do urban habitats induce physiological changes in Mediterranean lizards? J. Zool. 321, 75–82.

Voituron, Y., Roussel, D., Le Galliard, J. F., Dupoué, A., Romestaing, C. and Meylan, S. (2022). Mitochondrial oxidative phosphorylation response overrides glucocorticoid-induced stress in a reptile. J. Comp. Physiol. B Biochem. Syst. Environ. Physiol. 192, 765–774.

Wickham, H. (2016). ggplot2: elegant graphics for data analysis. New York: Springer.

Williams, R. (2019). The invasion ecology of common wall lizard (Podarcis muralis): population dynamics, interactions and adaptations.

Wist, B., Montero, B. K. and Dausmann, K. H. (2023). City comfort: weaker metabolic response to changes in ambient temperature in urban red squirrels. Sci. Rep. 13, 1–11.

Žagar, A., Simčič, T., Carretero, M. A. and Vrezec, A. (2015). The role of metabolism in understanding the altitudinal segregation pattern of two potentially interacting lizards. Comp. Biochem. Physiol. -Part A 179, 1–6.

